# Melanoma Addiction to GCDH is Mediated by NRF2 Tumor Suppressor Function

**DOI:** 10.1101/2021.10.22.465495

**Authors:** Sachin Verma, David Crawford, Ali Khateb, Yongmei Feng, Eduard Sergienko, Gaurav Pathria, Chen-Ting Ma, Steven H Olson, David Scott, Rabi Murad, Eytan Ruppin, Michael Jackson, Ze’ev A Ronai

**Author notes:** Correspondence: Ze’ev Ronai, Cancer Center, Sanford Burnham Prebys Medical Discovery Institute, 10901 N. Torrey Pines Rd, La Jolla, CA, 92037, USA. Email –.

## Abstract

Tumor dependency on specific metabolic signals has guided numerous therapeutic approaches. Here we identify melanoma addiction to the mitochondrial protein Glutaryl-CoA dehydrogenase (GCDH), a component in lysine metabolism which controls protein glutarylation. GCDH knockdown promoted apoptotic Unfolded Protein Response signaling and cell death in melanoma cells, an activity blocked by knockdown of the upstream lysine catabolism enzyme DHTKD1. Correspondingly, reduced GCDH expression correlated with improved survival of melanoma patients. A key mediator of GCDH-dependent melanoma cell death programs is the transcription factor NRF2, which induces ATF3, CHOP, and CHAC1 transcription linking lysine catabolism with the UPR signaling. NRF2 glutarylation upon GCDH KD increased its stability and DNA binding activity, which coincided with increased transcriptional activity, promoting apoptotic UPR signaling and tumor suppression. In vivo, genetic GCDH inhibition effectively inhibited melanoma tumor growth. Overall, these findings demonstrate an addiction of melanoma cells to GCDH, which by controlling NRF2 glutarylation limits apoptotic UPR signaling. Inhibiting the GCDH pathway could represent a novel therapeutic modality to treat melanoma.

## Introduction

Metabolic pathways that supply energy to normal cells^1, 2^ are often rewired in transformed cells to secure sufficient energy for rapid tumor cell proliferation^3, 4^. Among cancer relevant metabolic pathways are those functioning in uptake and utilization of amino acids, including glucose and glutamine, among other amino acids hubs^5, 6^. As a critical source of cellular wealth, amino acid catabolism is implicated in key homeostatic activities, including control of redox levels, ATP production, nucleotide biosynthesis and lipogenesis^5^. Common to these is tight control of fine-tuned signal transduction pathways, which are governed by spatial and temporal post translational modifications by various metabolites derived from amino acid catabolism (i.e., methylation, acetylation, malonylation, succinylation, and glutarylation)^5, 7^.

Addiction to particular metabolic pathways is common to tumor cells and often serves as their Achilles Heel in terms of vulnerability^8, 9^. Thus, there has been extensive effort to reverse tumor cell addiction by targeting a specific pathway or restricting availability of a particular amino acid^10^. Examples include strategies to limit glutamine, asparagine, serine/glycine or methionine; however, these approaches have met limited success, as sensing mechanisms promote compensatory growth factor or autophagic signaling^11–16^.

Although tumors have been the focus of these analyses, metabolic cues are also critical in the tumor microenvironment, including for anti-tumor immunity^17–19^. Thus, manipulating metabolic factors could be a double-edged sword: limiting availability of a particular amino acid to tumor cells may curtail an anti-tumor immune response, as reported in studies of glucose, glutamine and asparagine metabolism^20–23^. Combination therapies, in which metabolic signaling is blocked in the presence of other drugs that target oncogenic signaling^9^, have been proposed to overcome this hurdle. For example, limiting asparagine uptake while inhibiting MAPK signaling efficiently inhibits growth of pancreatic and melanoma tumor cells, with limited impact on immune cell function^14^. Likewise, targeting protein methyl transferase 5 (PRMT5) coupled with PD1 therapy is advantageous in targeting cold melanoma tumors^24^. Thus, it is critical to not only characterize metabolic pathways that underlie cancer cells’ ability to adapt to environmental or therapeutic pressure but also to identify their synthetic lethal partners^25^.

The essential amino acids lysine and tryptophan serve as building blocks for proteins and function in acetyl-CoA production and immunosuppression, activities critical for cancer cell survival^26, 27^. Lysine and tryptophan are degraded via a common pathway in which the dehydrogenase DHTKD1 catalyzes synthesis of the intermediate glutaryl CoA^5, 26, 28^. Glutaryl-CoA Dehydrogenase (GCDH) then converts glutaryl CoA to crotonyl CoA, which is metabolized to acetyl CoA to enter the TCA cycle^26, 27^. Interestingly, GCDH KO mice show elevated lysine glutarylation primarily in brain and liver, phenotypes suggesting that GCDH functions in TCA cycle-independent pathways^29, 30^ and that GCDH restricts lysine glutarylation by promoting glutaryl CoA breakdown. Notably, KO of genes encoding GCDH and DHTKD1 in mouse models^31, 32^ did not alter their viability, suggesting that the lysine catabolism pathway is dispensable for normal development and tissue homeostasis. However, when fed a high protein or high lysine diet, most GCDH KO mice die within a few days, pointing to the importance of GCDH for survival^33^ during protein catabolism. Lastly, coincident with GCDH loss and elevated lysine glutarylation^29^ is stabilization of the transcription factor NRF2^34^, a master regulator of the cellular stress response implicated in cellular oxidative, nutrient, UPR/ER and metabolic stress responses^35–38^. Enrichment of ER stress and unfolded protein response (UPR) genes was also observed as part of Keap1-mutant-specific vulnerabilities^39^ in lung adenocarcinoma tumor harboring hyperactive NRF2. Notably, NRF2 reportedly functions as both an oncogene^35–39^ and a tumor suppressor^40,41,42^, although mechanisms underlying the switch to either function are not well understood.

Here we characterize the importance of lysine and tryptophan catabolism for melanoma cells and identified their addiction for GCDH signaling. Furthermore, we show that GCDH activity controls NRF2 stability by regulating NRF2 glutarylation. GCDH loss promoted NRF2 glutarylation and increased its stability, promoting melanoma cell death via UPR signaling. Finally, we performed genetic inhibition of GCDH expression which suppress melanoma cell growth in culture and tumor growth *in vivo.*

## Results

### GCDH is required for melanoma cell survival

Lysine restriction have been shown to completely block cancer cell growth^43^ in colon carcinoma cell line. Likewise, the acetyl CoA generated from lysine catabolism was found to drive liver metastasis of colorectal cancer^27^.To directly assess the importance of the lysine catabolism pathway for melanoma cell viability we monitored the requirement of each of the component in this metabolic pathway for melanoma cell survival. To this end, siRNA inhibiting the expression of aminoadipic semialdehyde synthase (AASS), Kynurenine/alpha-aminoadipate aminotransferase (AADAT), Dehydrogenase E1 and transketolase domain containing 1 (DHTKD1), GCDH or Enoyl-CoA Hydratase, Short Chain 1 (ECHS1), components of the lysine catabolism pathway in A375 melanoma cells were used (Figure 1A). Surprisingly, only GCDH knock down (KD) resulted in a pronounced cell death in A375 cells (Figure 1B). These findings were confirmed using two independent siRNAs and recapitulated in two additional melanoma cell lines (UACC903, and 1205LU) (Figure 1C; Figure S1A and S1B). Correspondingly, GCDH KD upregulated cleaved caspase 3 and downregulated levels of the antiapoptotic markers BCL2/MCL1 in all three melanoma cell lines (Figure 1D; Figure S1C). Apoptotic signaling induced by GCDH KD was effectively reversed by treatment with the caspase inhibitor Emriscan (Figure 1E; Figure 2E). Notably, melanoma cell dependency on GCDH was found to be independent of mutational status (Figure 1F and G), indicating a metabolic signaling pathway that is uncoupled from BRAF or NRAS signaling pathways. These data points to the requirement of GCDH for melanoma cell survival.

**Figure 1.**
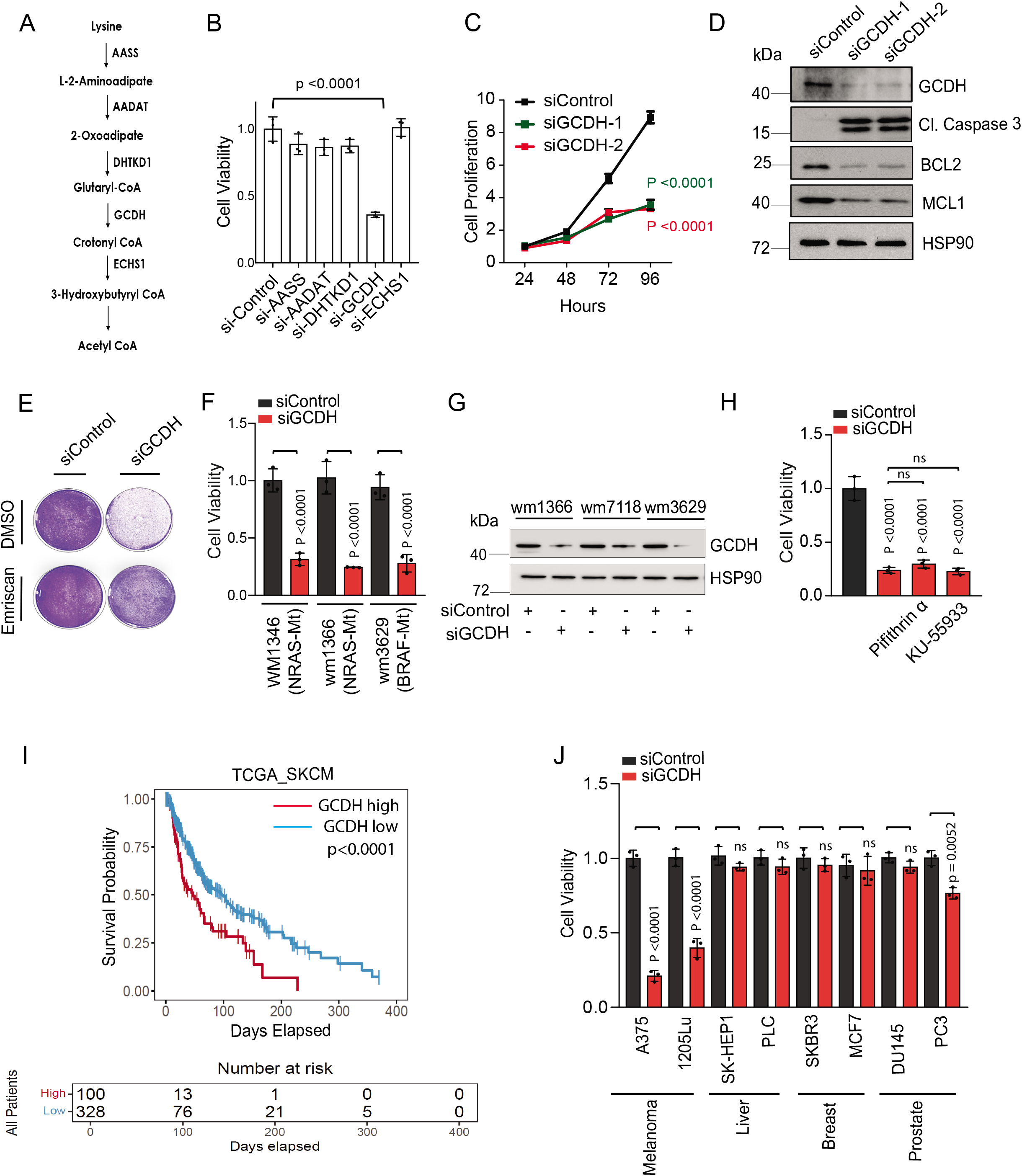
GCDH is required for melanoma cell survival. (A) Schematic representation of enzymes involved in lysine catabolic pathway. (B) siRNA targeting AASS, AADAT, DHTKD1, GCDH, ECHS1 or control sequence were transfected into A375 melanoma cells by Jetprime for 96 hours. Cell viability was then measured by quantifying crystal violet staining. (C) Cell growth upon GCDH knock down using two independent siRNAs for 0-96 hr in A375 cell line. Cell growth was analyzed by cell counting at indicated time points. (D) A375 cells were transfected with siRNA against GCDH, and western blot analysis was done using indicated antibodies. (E) Cell viability assay of control or GCDH KD A375 cells treated with the caspase inhibitor Emriscan (10μM for 48 hr). (F) Cell viability assay of indicated cells 96 hr after transfection with siRNAs targeting GCDH. Cell viability was measured by quantifying crystal violet staining. (G) Western blot analysis confirming GCDH KD as described in D. (H) Analysis of cell viability upon GCDH-KD alone or in combination with treatment with Pifithrin-α or KU-55933 in A375 cells. (I) Survival analysis of GCDH expression in melanoma patients using TCGA. Number of total patients, n =428, n (GCDH high) =328 and n (GCDH low) =100. (J) Cell viability was measured by quantifying crystal violet staining upon GCDH KD in various cancer cell lines as indicated. (K) Western blot analysis on A375 cells transfected with siRNA targeting GCDH for 96 hours using indicated antibodies. Data are represented as mean ± SEM of n = 3 independent experiments. Statistical significance (indicated p value or ns-not significant w.r.t control) was calculated using unpaired *t*-test.

Cell death seen upon GCDH KD in A375 melanoma cells (p53 wt) could not be rescued upon pharmacological inhibition of either p53 (Pifithrin-α) or ATM (KU-55933) nor was it affected upon treatment with the ROS scavenger N-acetylcysteine (Figure 1H and 3E). These findings imply that melanoma dependency on GCDH is independent of DNA damage or oxidative stress. We also asked whether GCDH inhibition altered mitochondrial biogenesis or cellular respiration. To do so, we performed GCDH KD and measured basal, maximum oxygen consumption rate (OCR) and spare respiratory capacity. None of these were altered significantly in GCDH KD versus control cells (Figure S1D), indicating a lack of effect on mitochondrial biogenesis and respiration. Notably, neither GCDH nor any other component of the lysine catabolism pathway was required for cell viability in the non-transformed and immortalized melanocyte cell line H3A (Figure S1E). This observation is consistent with GCDH and DHTKD1-KO mouse phenotype where GCDH activity was found to be dispensable for overall viability and growth of normal cells.

Evaluation of melanoma patient samples revealed that higher GCDH expression was associated with decreased patient survival (Figure 1I), whereas patients with low levels of GCDH expression exhibited a significant survival advantage, compared with those with high levels. Notably, such correlation was not seen in patients with other tumor types (Figure S2).

Consistent with patient data, inhibiting GCDH expression in melanoma lines, but not in liver, breast or prostate cancer cultures, promoted notable cell death (Figure 1J and S1F), highlighting the specificity of GCDH signaling in melanoma. These data highlight the importance of GCDH for melanoma cell survival.

### GCDH inhibition promotes UPR-dependent cell death signaling

To identify possible mechanisms that underlie cell death induced upon GCDH loss, we monitored changes in gene expression following GCDH KD in melanoma cells. RNAseq analysis followed by IPA assessment of signaling pathways that were differentially expressed upon GCDH KD identified UPR, sirtuin, GADD45, pentose phosphate, p53 and ATM signaling pathways (Figure S3A). Among differentially expressed genes were upregulation of genes controlled by ATF3 and ATF4 signaling implicated in UPR-induced cell death (i.e., DDIT3, -CHAC1, and GADD45a)^44^, and genes implicated in cell cycle inhibition and tumor suppression (i.e., CDKN1A and CDKN2B^45–47^ (Figure 2A and B). qPCR analysis of gene signatures performed in both A375 and UACC903 cells confirmed upregulation of apoptotic UPR signaling as reflected by increased levels of ATF3, ATF4, CHOP and CHAC1 transcripts upon GCDH KD (Figure 2C and S3B) in A375 and UACC903 cells respectively. Inhibition of either ATF3, DDIT3 or CHAC1 in A375 melanoma cells subjected to GCDH KD, effectively attenuated the degree of cell death induced upon GCDH KD (Figure 2D). These findings overall suggest that GCDH loss in melanoma cells induces UPR-dependent cell death pathways regulated by the ATF3-DDIT3-CHAC1 apoptotic cascade^44^.

**Figure 2.**
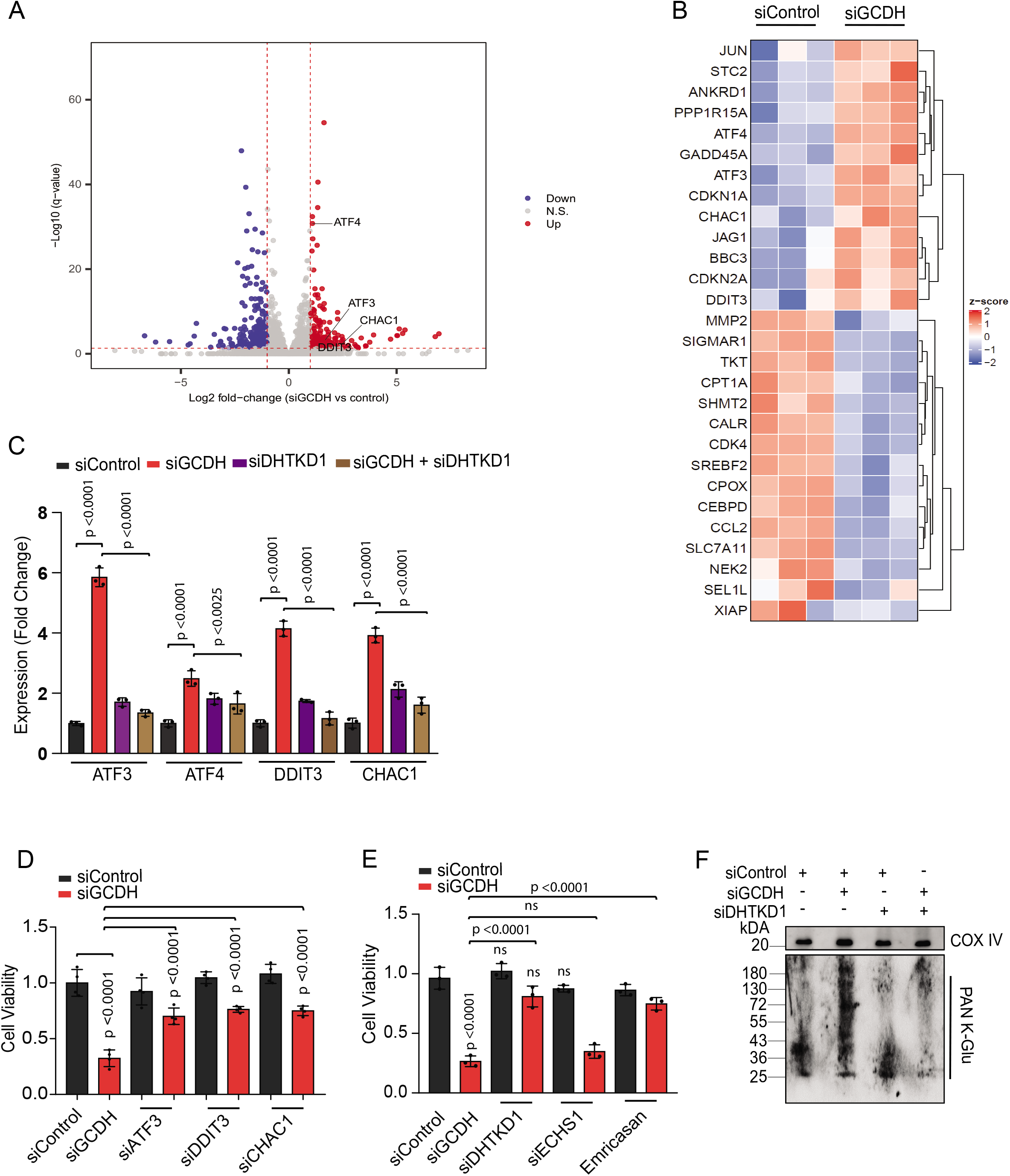
GCDH inhibition promotes UPR-dependent cell death signaling. (A) Volcano plot showing elevated expression of molecules controlling UPR mediated cell death cascade (ATF3, ATF4, DDIT3 and CHAC1), identified by RNA-seq analysis. (B) Heatmap representing differential expression of ATF3/4 downstream targets in GCDH-KD A375 cells identified by RNA-seq analysis. (C) RT_qPCR validation of ATF3, ATF4, DDIT3 and CHAC1 in A375 cells transfected with indicated siRNAs. (D and E) Cell viability of A375 cells transfected with indicated siRNAs for 96 hours. Cell viability was analyzed by crystal violet staining and quantitation. (F) Western blot analysis on mitochondrial extracts from A375 cells using PAN K-Glu-detecting antibody to detect lysine glutarylation 72 hr following transfection with indicated siRNAs. Data are presented as the mean ± SEM. Statistical significance (indicated p value or ns-not significant w.r.t control) was calculated using unpaired *t*-test and two-way ANOVA for (C) and (E).

Consistent with apoptotic signaling observed at molecular level (Figure 1D and S1C), GCDH KD in A375 cells led to a 50.1 % increase in cells exhibiting DNA fragmentation, indicative of cell death, relative to controls (4.4 %) (Figure S3C). Significantly, the degree of cell death seen upon GCDH KD was decreased to 14.1 % by concomitant KD of DHTKD1, which is upstream of GCDH in the pathway (Figure 1A, S3C and S3D). While DHTKD1 KD also effectively rescued viability of BRAF mutant 1205LU melanoma cells subjected to GCDH KD, (Figure S3E), KD of ECHS1, which is downstream of GCDH (Figure 1A), did not alter the degree of cell death seen in the presence of GCDH KD (Figure 2E). Given that GCDH inhibition promotes glutaryl-CoA accumulation and increased protein glutarylation in mitochondria^29^, we assessed protein glutarylation in GCDH KD cells. Relative to control siRNAs, GCDH KD promoted increased glutarylation of proteins in mitochondrial extracts (Figure 2F). Accordingly, levels of glutarate, a metabolite produced by glutaryl-CoA, increased in GCDH KD A375 cells (Figure S3F). Changes seen in both glutarylation and glutarate levels were largely rescued upon DHTKD1 KD (Figure 2F and S3F). Levels of ATF3, ATF4, DDIT3 and CHAC1 transcripts were also attenuated upon combined KD of DHTKD1 and GCDH (Figure 2C), rescuing changes seen upon GCDH KD alone. These observations suggest that GCDH controls levels of protein glutarylation, which in turn regulate UPR cell death signaling.

### GCDH loss in melanoma cells increases NRF2 levels and induces NRF2 dependent UPR cell death

Next, we asked whether NRF2 function is required to induce apoptotic UPR signaling after GCDH KD, given that NRF2 is upregulated in the striatum of GCDH KO mice with high lysine diet and Quinolinic acid induced toxicity^34^ and implicated in the regulation of ATF3 and ATF4 transcription^37, 48, 49^. Since NRF2 is regulated at the transcriptional level and protein stability, we monitored potential changes in NRF2 protein and transcript levels after GCDH KD in A375 and UACC903 lines (Figure 3A, S4A and S4B). We found that NRF2 mRNA level were only marginally affected upon GCDH KD (Figure S4A). Whereas elevated NRF2 protein levels coincided with increased abundance of the UPR proteins ATF3, ATF4, DDIT3, CHAC1, caspase 3 (cleaved) and the downstream NRF2 targets HO1 and p21 (Figure 3A and S4B). Concomitant KD of NRF2 or DHTKD1 in GCDH KD A375 and UACC903 lines effectively reversed apoptotic UPR signaling seen upon GCDH KD alone, both at the protein (Figure 3A and S4B) and transcript (Figure 3B and Figure S4C) levels. Given that ATF3 controls DDIT3-CHAC1^37, 44^ signaling, we asked whether ATF3 may mediate phenotypes seen after GCDH loss. ATF3 KD combined with GCDH KD in these lines effectively attenuated cell death seen in the presence of GCDH KD alone (Figure 2D), similar to the effects seen following DHTKD1 KD (Figure 2E). ATF3 KD in GCDH KD cells was also accompanied by reduced levels of ATF4, DDIT3, CHAC1 and cleaved caspase 3 protein (Figure 3C and S4D). Consistently, decreased levels of ATF4, DDIT3 and CHAC1 transcripts, was observed in cells subjected to ATF3 KD, compared with GCDH KD (Figure 3D and S4E). The changes seen upon ATF3 KD phenocopied those observed following DHTKD1 KD, or NRF2 KD in cells that were subjected to GCDH KD, culminating in attenuated apoptotic UPR signaling (Figure 2D and 3E). As expected, ATF3 KD alone in melanoma cells had no effect on NRF2 stability or p21 or HO1 expression, suggesting that ATF3 is the primary driver of apoptosis downstream of NRF2 after GCDH inhibition (Figure 3C and S4D). Notably, NRF2 KD in the presence of GCDH KD decreased the extent of melanoma cell death seen after GCDH KD alone (Figure 3E). Collectively, these observations suggest that GCDH controls apoptotic UPR signaling in melanoma cells via elevated abundance of NRF2protein levels.

**Figure 3.**
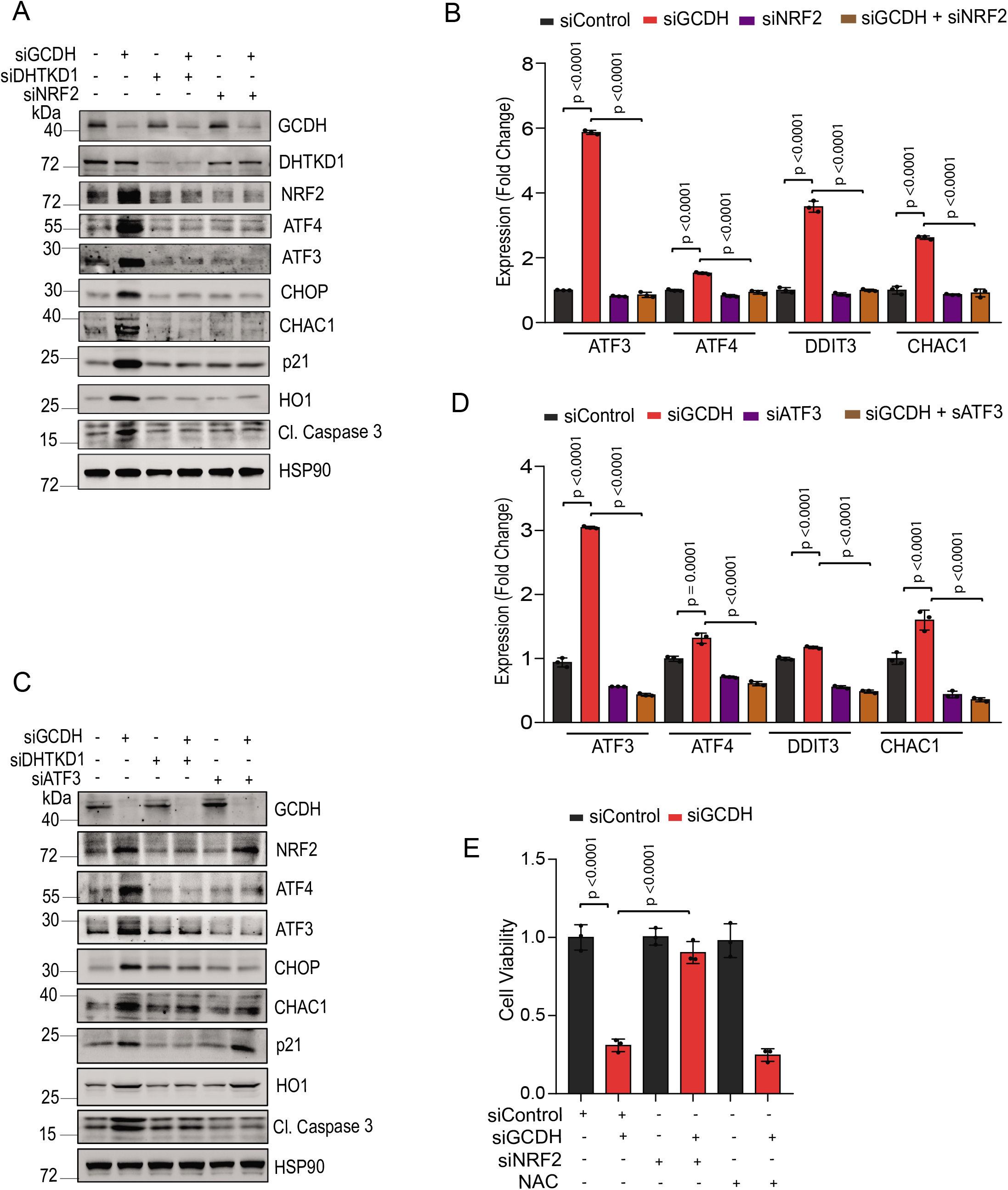
GCDH loss in melanoma cells increases NRF2 levels and enhances UPR/cell death signaling. (A) Western blot analysis of indicated proteins in A375 cells 72 hr following transfection with indicated siRNAs. (B) RT-qPCR analysis of *ATF3, ATF4, DDIT3,* and *CHAC1* expression levels in A375 cells following transfection with indicated siRNAs. (C) Western blot analysis of indicated proteins in A375 cells 72 hr following transfection with indicated siRNAs. (D) RT-qPCR analysis of *ATF3, ATF4, DDIT3,* and *CHAC1* expression levels in A375 following transfection with indicated siRNAs. (E) Viability assay on A375 cells upon transfected with indicated siRNAs and treated or untreated with NAC (10mM) for 48 hr. Data are presented as the mean ± SEM. Statistical significance (indicated p value or ns-not significant w.r.t control) was calculated using two-way ANOVA for (B), (D) and (E).

### GCDH controls NRF2 stability

We next monitored potential changes in NRF2 stability following GCDH KD in A375 cells. NRF2 half-life increased following GCDH KD in A375 melanoma cells relative to controls (Figure 4A and 4B) as well as in GCDH KD HEK293T cells exogenously expressing HA-NRF2 (Figure S5B and S5A). Conversely, DHTKD1 KD decreased NRF2 stability (Figure S5C and S5A). As NRF2 stability is tightly controlled by interaction with the ubiquitin ligase KEAP1^50^, we assessed possible changes in NRF2/KEAP1 interaction following GCDH KD. Immunoprecipitation (IP) of endogenous NRF2 from GCDH KD cells revealed lower levels of NRF2-bound KEAP1 relative to control cells (Figure 4C). Given notable increases in protein glutarylation seen upon GCDH inhibition, we asked whether NRF2 glutarylation altered its interaction with KEAP1 or degree of ubiquitination, given that lysine is the primary glutarylated residue. To do so, we subjected NRF2 immunoprecipitates from melanoma cells to immunoblotting with antibodies against lysine glutarylation (K-Glu). While basal levels of NRF2 glutarylation were detected in control A375 cells, those levels notably increased following GCDH KD (Figure 4C). Likewise, IP of ectopically expressed HA-NRF2 in HEK293T cells followed by immunoblotting with K-Glu antibodies revealed elevated NRF2 glutarylation, compared to controls (Figure S5D). To confirm NRF2 glutarylation, we performed an *in vitro* glutarylation assay with purified HA-NRF2 and observed its glutarylation (K-Glu NRF2; Figure 4D) when incubated with glutaryl-CoA harboring a reactive CoA moiety but not with glutaric acid, which served as control, suggesting that NRF2 undergoes glutarylation in the presence of elevated glutaryl-CoA levels promoted by GCDH KD. We next examined relative amounts of KEAP1-bound to glutarylated NRF2. IP of HA-NRF2 or K-Glu HA-NRF2 (used as bait) to retain KEAP1 from extracts of A375 cells showed lower interaction of glutarylated NRF2 with KEAP1 (Figure 4E). Moreover, cell fractionation revealed that glutarylated NRF2 was primarily nuclear (Figure S5E), implying effects on transcriptional activity. Accordingly, we performed an electrophoretic mobility shift assay to monitor changes in NRF2 binding to the antioxidant response element (ARE), a promoter element in genes which is bound and regulated by NRF2. Relative to the HA-NRF2 control, we observed a notable increase in binding of *in vitro* glutarylated K Glu HA-NRF2 to the ARE (Figure S5G), suggesting increased NRF2 affinity for the promoter sequence. RNAseq data performed in melanoma cells after GCDH inhibition confirmed a gene expression signature (Figure 2A and S3A) different from that seen in control cells^36^ but consistent with an NRF2-activated UPR signature^38^. These observations suggest that GCDH control of NRF2 glutarylation not only determines its stability but also modulates its binding to DNA.

**Figure 4.**
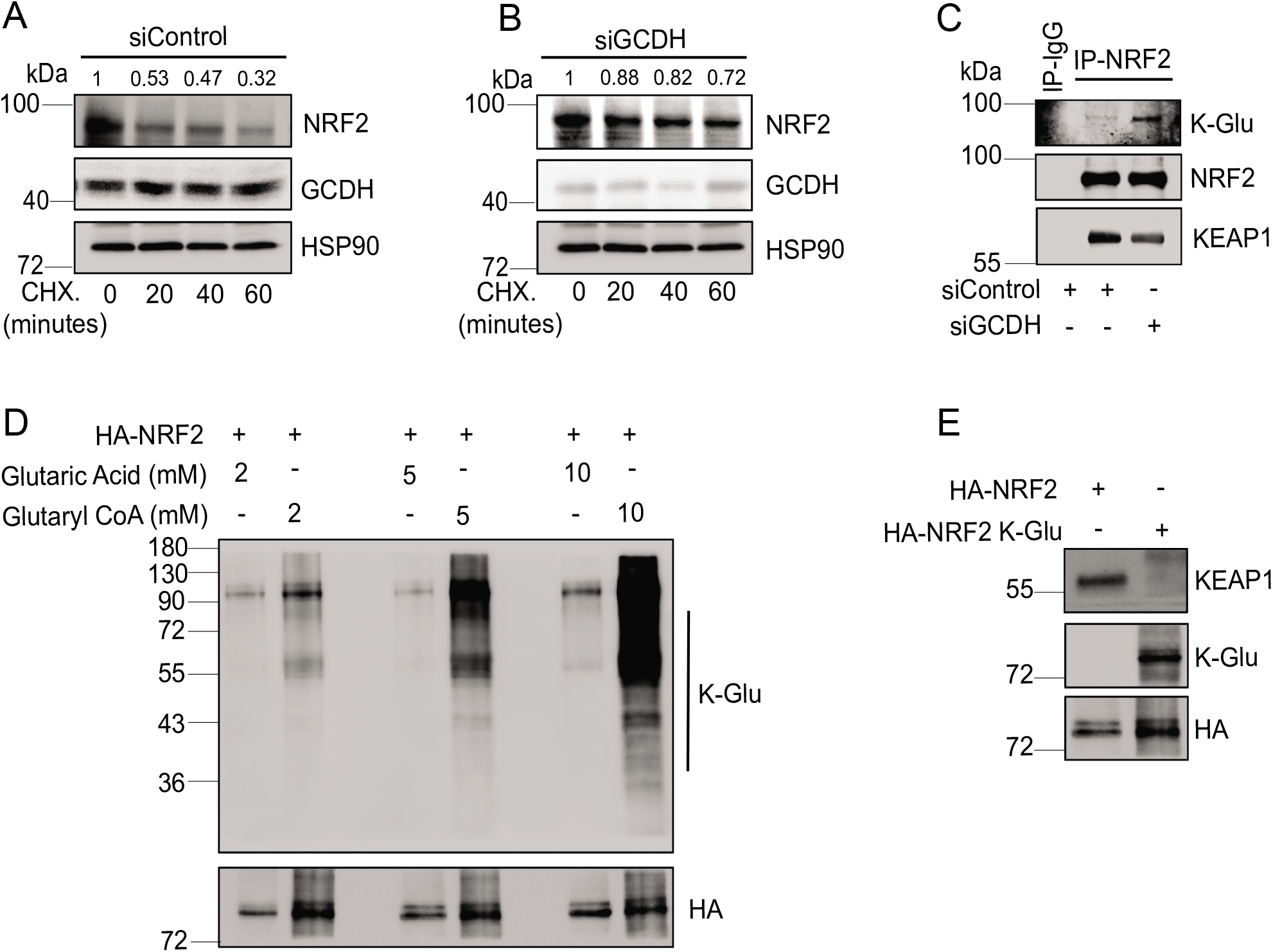
Lysine glutarylation increases NRF2 stability by attenuating KEAP1 binding. (A, B) Cycloheximide (CHX) chase to check half-life of endogenous NRF2 in A375 cells transfected with siRNA targeting GCDH. Western blot was performed on lysate from A375 transfected with siControl (A) or siGCDH (B) for 72 hours and then treated with 10 μM Cycloheximide (CHX) for indicated time. After quantification, the signals obtained in panel A and B were used to calculate the NRF2/HSP90 ratios and described with respect to CHX. treatment period. (C) Immunoprecipitation and Western blot analysis of A375 transfected with indicated constructs. Cells were treated with the proteasomal inhibitor MG132 for 4 hr followed by IP/Western blotting analysis with antibodies to detect K-Glu PTM and NRF2. (D) In vitro glutarylation assay on purified HA-NRF2 following incubation with indicated concentration of glutaryl CoA. (E) In vitro KEAP1 binding analysis performed using purified HA-NRF2 or K-Glu-NRF2 as bait on A375 cell lysates. A representative image of n = 3 independent experiments is shown.

### The GCDH inhibition suppress melanoma growth in vivo

We next monitored effects of genetic GCDH inhibition on melanoma growth *in vivo* using immunodeficient mice inoculated with human A375 melanoma cells. Genetic inactivation of GCDH was achieved by cloning shRNA targeting GCDH under doxycycline inducible promoter in A375 melanoma cells, that were then inoculated into nude mice. To induce genetic inhibition of GCDH mice were fed with DOX containing chow. Indeed, inhibition of GCDH expression in vivo lead to attenuation of tumor growth, compared to control GCDH expressing tumors (Figure 5A). Inhibition of GCDH expression and activation of NRF2 and concomitant downstream UPR cell death markers was confirmed for ATF3, CHAC1 and cleave caspase 3 in each of the 7 tumor lysates from GCDH KD compared with control experimental groups (Figure 5B). These finding substantiate the importance of GCDH for melanoma growth; inhibition of GCDH expression results in effective melanoma growth inhibition *in vivo*.

**Figure 5.**
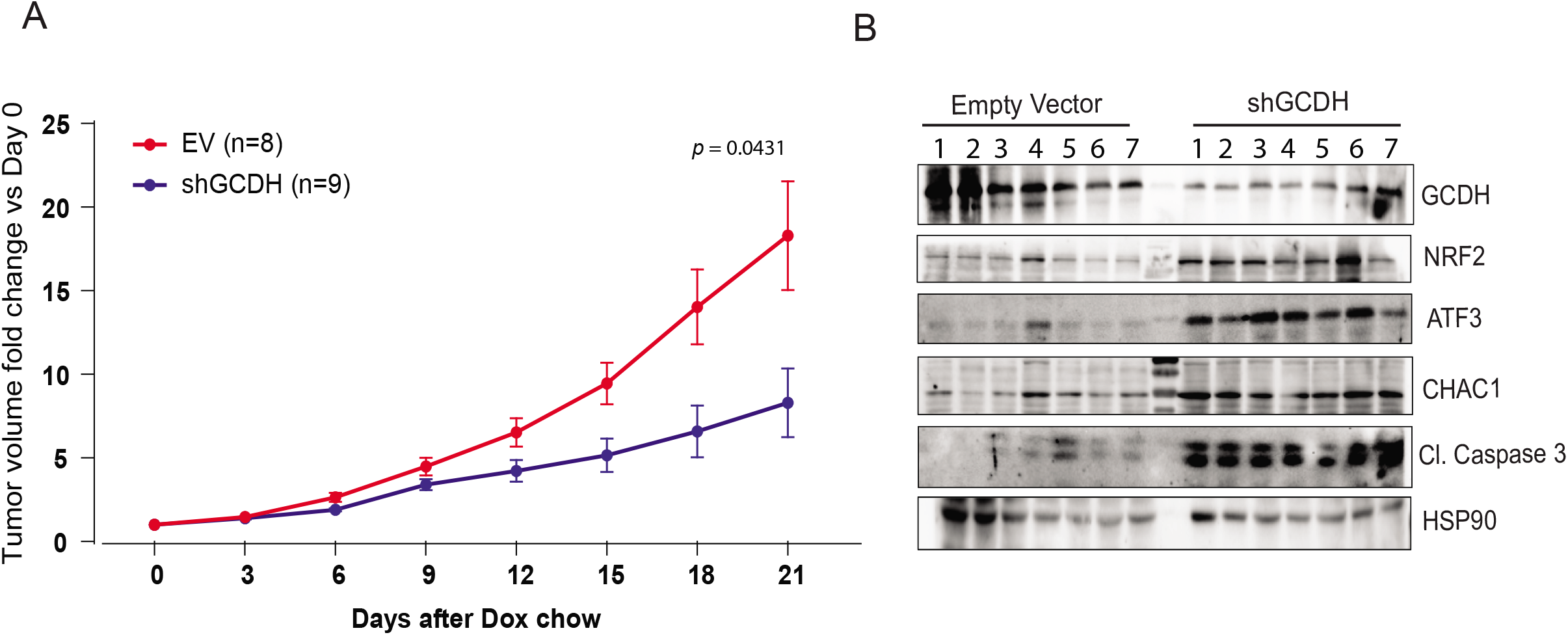
GCDH inhibition using inducible shRNA attenuates melanoma proliferation and tumorigenesis. (A) Fold change in tumor volume of human melanoma A375 cell line following dox chow treatment. NOD/SCID (NOD.CB17-Prkdcscid/J) mice were injected subcutaneously with 1 ×10^6^ A375 cells. (B) Western blot analysis present protein levels of GCDH, NRF2, ATF3, CHAC1 and Cl. Caspase 3 in tumor harvested from tumors subjected to control or GCDH KD detailed in panel A. Data are presented as the mean ± SEM. Statistical significance (indicated p value relative to control) was calculated using paired *t*-test.

## Discussion

Extensive efforts have been made to identify tumor cell vulnerability to changes in metabolic signaling, with the goal of curtailing selective metabolic cues to limit tumor but not normal cell growth. Nonetheless, the number of drugs that have reached clinical evaluation remains limited. This is because tumor cells manage to activate alternate metabolic pathways to compensate for attenuated metabolic flux, or because the targeted metabolic pathway impairs normal cell function and/or curtails the immune response or other microenvironmental factors that limit tumor growth. Here we establish a novel paradigm and demonstrate melanoma cell addiction to GCDH, one of the enzymes in the multi-step lysine catabolism pathway. Moreover, blocking GCDH activity, but not that of upstream or downstream components of the lysine catabolism pathway, resulted in significant tumor cell death. Our studies define NRF2 as the principal component mediating apoptotic UPR signaling that induces cell death programs. NRF2 glutarylation, seen following GCDH KD, stabilizes NRF2 and likely enhances its transcriptional activation of factors mediating apoptotic UPR signaling. Knockdown of either NRF2, its pro-apoptotic transcriptional target ATF3^37, 40^ or GCDH-upstream enzyme DTHKD1 effectively blocked cell death phenotypes seen upon GCDH KD, indicating that a pathway controlled by GCDH activity allows survival of melanoma cells.

NRF2 reportedly exhibits both oncogenic^35,38^ and tumor suppressor activities^40,41,42^, in different cancer models, although mechanisms determining those activities are not well understood. In melanoma, NRF2 has been previously shown to affect innate immune responses and oxidative stress^51^. Additionally, high levels of NRF2 protein were found to be associated with a poor prognosis in melanoma irrespective of oxidative stress^52^. Our findings show that NRF2 exhibits tumor suppressor activity upon glutarylation and suggest that NRF2 glutarylation induced by GCDH loss-of-function both promotes its dissociation from the E3 ubiquitin ligase KEAP1 and enhances it stability, which then increases NRF2-dependent expression of select gene set that mediate apoptotic UPR signaling. NRF2 glutarylation occurs on lysines that may otherwise serve as ubiquitin acceptor sites, reducing its ubiquitination and enhancing its stability. Mapping NRF2 lysine glutarylation site(s) would be desirable; however, similar to lysine ubiquitination, glutarylation may be promiscuous, such that when some sites are unavailable, others are modified. Relevant to transcriptional effects, we perform in vitro gel shift assays and showed that NRF2 glutarylation enhances its binding to the known NRF2 response element. However, it is also possible that glutarylated NRF2 possesses greater affinity to form complexes with transcriptional co-activators or co-suppressors, or with epigenetic regulators governing translation initiation complex assembly, each of which would define a select transcriptional readout.

Our findings demonstrate that addiction to GCDH signaling is observed in melanoma cells but not in breast, colon or prostate tumor cells. Important support for these findings comes from clinical data, in which low GCDH expression coincided with better patient outcomes in melanoma but not in other tumor types. One explanation for selective GCDH dependency is the neural crest origin of melanoma^53^, resembling phenotypes seen in brain of the GCDH KO mice^31^. Equally plausible is that different proteins undergo glutarylation following GCDH loss in tumors other than melanoma, which may alter different signaling pathways. In support of this possibility is the observation of numerous glutarylated proteins upon GCDH KD^29,30^ in different tissues, which could modify activity of distinct drivers of oncogenesis depending on the substrate

Would targeting GCDH offer a novel therapeutic modality for melanoma? Data from total KO mice suggest that ablation of either GCDH or other components of the lysine catabolism pathway^31, 32^ does not have a major impact on either normal development or tissue homeostasis, and mice are viable with minor deficiencies. However, mice globally deficient in GCDH acquire vulnerability to excessive lysine or high protein diets^33^, implying that a ketogenic diet may enhance cell death in GCDH-low tumor cells, a possibility deserving further assessment. Our in vivo data supports effectiveness of genetic GCDH inhibition, which attenuated melanoma growth in immunodeficient mice, suggesting that GCDH may be required for tumor cell growth *in vivo.* Further work is required to examine the effect of melanoma addiction to GCDH on the TME including anti-tumor immunity. Melanoma addiction to GCDH well illustrates the selective advantage of select metabolic cue for distinct tumor types.

## Methods

### Animal studies

All animal procedures were approved by the Institutional Animal Care and Use Committee (IACUC) of Sanford Burnham Prebys Medical Discovery Institute (approval number AUF 19-082). Animal experiments were performed at Sanford Burnham Prebys Medical Discovery Institute Animal Facility in compliance with the IACUC guidelines. The study is performed with all relevant ethical regulations regarding animal research. The xenograft model was established using A375 cells expressing either control or shRNA targeting GCDH using doxycycline inducible PLKO-1 vector. NOD/SCID (NOD.CB17-Prkdcscid/J) mice were obtained from the SBP Animal Facility. Eight-week-old male C57BL/6 mice were injected subcutaneously in the flank with 1 × 10^6^ A375 cells. For in vivo GCDH KD experiments, mice were fed rodent chow containing 200 mg/kg doxycycline (Dox diet, Envigo) to induce GCDH-KD. Mice were sacrificed upon signs of morbidity resulting from tumor growth.

Tumor volume was measured with linear calipers and calculated using the formula: (length in mm × width in mm) x 1/2. After the mice were sacrificed, tumors were frozen or fixed in Z-Fix (Anatech). Snap-frozen tumors were utilized for protein extraction for further analysis.

### Cell culture and reagents

Cancer cell lines (breast: SKBR3 and MCF7; prostate: PC-3 and DU145; liver: SK-HEP1 and PLC and melanoma: A375 and 1205LU) were obtained from ATCC. WM1346 and WM1366 and WM3629 melanoma cell lines were a gift from M. Herlyn (Wistar Institute) and UACC-903 melanoma cell line was obtained from the University of Arizona Cancer Center. All cell lines were cultured in Dulbecco’s modified Eagle’s medium (DMEM; GE Healthcare), supplemented with 5% fetal bovine serum and penicillin-streptomycin. All cells were grown at 37 °C in a humidified atmosphere containing 5% carbon dioxide. Amino acids (L-lysine, L-arginine), proteasomal inhibitor (MG132), ATM Inhibitor (KU-55933) and p53 inhibitor (Pifithrin-α) were purchased from Sigma-Aldrich.

### siRNA, DNA constructs, transfection and transduction

1×10^5^ cells were seeded overnight (O/N) per well in 6-well plates. Negative control (NT-siRNA) or si-RNA targeting the transcript of interest was transfected utilizing jetPRIME® transfection reagent, as per manufacturer’s instructions (Polyplus, NY, USA). Following siRNAs were used: si-GCDH (SASI_Hs01_00246318 and SASI_Hs01_00246319), siDHTKD1 (SASI_Hs02_00352234 and SASI_Hs02_00352235), si-ATF4 (SASI_Hs02_00332313), si-ATF3 (NM_001030287), si-DDIT3 (SASI_Hs01_00153013), si-CHAC1 (SASI_Hs01_00146246), si-NRF2 (SASI_Hs01_00182393), si-AASS (SASI_Hs02_00340239 and SASI_Hs01_00016127) si-ADAT (SASI_Hs01_00091485 and SASI_Hs01_00091486) siECHS1 (SASI_Hs01_00085563 and SASI_Hs02_00336896) and the negative control si-RNA (NT-siRNA; SIC001) were purchased from Sigma-Aldrich. Plasmid encoding HA-NRF2 (pLV-mCherry-CMV HA-NRF2) was synthesized by Vector builder, USA).

### Cell proliferation and viability

For cell proliferation, 0.3-0.5×105 cells were seeded O/N in triplicate in 6-well plate. Following treatment for the specified duration, cells were trypsinized and the cell count was determined with Neubauer hemocytometer (Celeromics, Cambridge, UK). To measure cell viability, cells were washed twice with cold PBS and fix for 10 minutes with ice-cold 100% methanol. Cells were then incubated with 0.5% crystal violet solution in 25% methanol for 10 minutes at room temperature. Crystal violet solution was removed, and cells washed in water several times, until the dye stops coming off. The culture plates were then dry at room temperature. For quantitation, 2 ml 10% acetic acid to each well (6 well) was incubate for 20 min with shaking (extraction). 0.5-1 ml of extracted stain was diluted 1:4 in water (dilution flexible based on signal intensity) followed by quantifying absorbance at 590 nm. The resulting reading were normalized as compared to absorbance from respective control cells.

### SubG1 DNA content analysis to quantify apoptotic population

SubG1 DNA content analysis was performed for determination of apoptotic population, analyzed by propidium iodide staining (Sigma Aldrich). Briefly, 1 × 106 cells were washed twice with cold PBS and fixed in 70% ethanol in PBS at 4°C overnight. Cells were washed, pelleted by centrifugation, and treated with RNase A (100 μg/mL) and propidium iodide (40 μg/mL) at room temperature for 30 min. Cell cycle distribution was assessed by flow cytometry (BD LSRFortessa™, BD Biosciences), and data was analyzed using FlowJo software.

### Immunoblotting

Total protein was extracted in Laemmli buffer, fractionated by SDS polyacrylamide gels and transferred to PVDF membranes (Millipore Sigma, MA, USA). After blocking with 5% non-fat dry milk (BD Biosciences, CA, USA), the membranes were incubated with primary antibodies overnight at 4°C. Afterwards, 2 hr incubation with HRP – conjugated secondary antibodies was performed. Following chemiluminescence reaction, the protein signal was visualized using the ChemiDoc imaging system (Bio-Rad, Hercules, CA) according to the manufacturer’s instructions.

### qPCR Analysis

1×105 cells were seeded O/N per well in 6-well plates. Following treatment for 72 hr, total RNA was extracted using RNeasy Mini Kit (Qiagen, Hilden, Germany). cDNA was synthesized using oligo(dT) and random primers (AB Bioscience, MA, USA), and qPCR analysis was performed with SYBR Green (Roche, NJ, USA). Primers were designed using the PrimerQuest tool (Integrated DNA Technologies, CA, USA) and Primer Bank (https://pga.mgh.harvard.edu/primerbank/). Œl-Actin was used as an internal control. Primer efficiency was measured in preliminary experiments, and amplification specificity was confirmed by dissociation curve analysis.

Sequence of the primers used:

ATF3-F: CCTCTGCGCTGGAATCAGTC

ATF3-R: TTCTTTCTCGTCGCCTCTTTTT

ATF4-F: GTATGAGCCCAGAGTCCTATCT

ATF4-R: CACATTGACGCTCCTGACTATC

CHAC1-F GAACCCTGGTTACCTGGGC

CHAC1-R CGCAGCAAGTATTCAAGGTTGT

DDIT3-F GGAAACAGAGTGGTCATTCCC

DDIT3-R CTGCTTGAGCCGTTCATTCTC

GCDH-F: CGTCCCGAGTTTGACTGGC

GCDH-R: GATGCGAGGCATGAGTCTCT

β-Actin-F: CATGTACGTTGCTATCCAGGC

β-Actin-R: CTCCTTAATGTCACGCACGAT

### Intracellular Glutarate Quantification

Cell extraction and GC-MS analysis for Glutarate quantification was performed as described in^54^. Intracellular metabolite amounts are expressed in nmol per cell sample (cells from one well of six-well plates; approximately 0.5 × 106-1.0 × 106 cells).

### Antibodies

The following antibodies were used: NRF2 (D1Z9C) (dilution, 1:1,000), ATF4 (D4B8) (dilution, 1:1,000), CHOP (L63F7) (dilution 1:1,000), p21 Waf1/Cip1 (12D1) (dilution 1:1,000), HO1 (E9H3A) (dilution, 1:1,000), Cleaved Caspase-3 (Asp175) (5A1E) (dilution 1:1,000) and HRP-conjugated anti-Mouse (dilution 1:10,000) and antiRabbit (dilution, 1:10,000) antibodies from Cell Signaling Technology (MA, USA). Bcl-2 (C-2) (dilution 1:1,000) Mcl-1 Antibody (22) (dilution, 1: 1,000) and HSP90 (F-8) (dilution, 1: 5,000) from Santa Cruz Biotechnology, (TX, USA). GCDH/GCD antibody (ab232774) (dilution 1:10,000) DHTKD1 antibody (ab230392) (dilution 1:10,000) antibodies from Abcam. Pan Anti-glutaryllysine antibody from PTM Biolab LLC. CHAC1 (dilution 1: 5 00) antibody from Proteintech.

### Cycloheximide chase assay

Cycloheximide chase was performed as previously described. Briefly, cycloheximide (50 μg/ml) was added to cells for indicated times, and cell lysates were analyzed with indicated antibodies.

### Immunoprecipitation

The protein–protein interaction for endogenous protein was studied by using the Co-Immunoprecipitation Kit (Thermo Scientific). For overexpressed HA-tag proteins, HEK293T cells were transfected with plasmids encoding the HA-NRF2 (pLV-mCherry-CMV HA-NRF2, Vector builder, USA). After 48 hours of transfection, cells were treated with MG132 for 4 hours and washed with PBS. Cells were lysed in IP milder Lysis/Wash Buffer (0.025M Tris, 0.15M NaCl, 0.001M EDTA, 1% NP-40, 5% glycerol; pH 7.4) and HA-antibody-conjugated agarose resin (Life technology) was added. It was rotated overnight at 4 °C. After incubation, resin was pelleted and washed with IP Lysis/Wash Buffer. It was boiled in 2 × SDS-PAGE loading buffer for 5 min and analysed by western blotting. For detection of glutarylated NRF2 from cell fractions (MF-membrane, NF-nuclear and CF-cytoplasmic), first cell fractionation was carried out using Cell Fractionation Kit (CST #9038). Equal amount of protein from purified fractions were incubated with HA beads overnight at 4° C.

### *In vitro* glutarylation, KEAP1 binding and ARE-EMSA

HEK293T cells expressing HA-NRF2 were lysed in RIPA buffer (20mM Tris-HCl (pH 7.5), 150 mM NaCl, 1 mM Na2 EDTA, 1 mM EGTA, 1% NP-40, 1% sodium deoxycholate, 2.5 mM sodium pyrophosphate, 1 mM b-glycerophosphate, 1 mM Na3 VO4, 1 μg/ml leupeptin) containing N-Ethylmaleimide (NEM). The lysates were incubated with HA binding beads and washed with RIPA lysis buffer to retain HA-NF2 on beads and subjected to in vitro glutarylation reaction. For in Vitro Glutarylation of NRF2, purified HA-NRF2 in glutarylation buffer (50 mM HEPES [pH 8.0], 150 mM NaCl, and protease inhibitors) were mixed with different concentration of glutaryl-CoA or gluataric acid to form glutaryl-NRF2 (K Glu-NRF2) and control HA-NRF2 respectively. The reactions were incubated in an Eppendorf Thermomixer for 4 hr at 37°C at 400 rpm. To minimize condensation, samples were briefly centrifuged every hour during incubation. The washed HA-beads (described above) were then used for KEAP1 binding or EMSA directly. For KEAP1 binding assay, beads were incubated with cell extract (for KEAP1 binding) in IP lysis/wash buffer. After subsequent washing steps beads were boiled in 2X SDS sample buffer, the proteins were separated on SDS/PAGE and immunoblotted with indicated antibodies as described. For ARE-EMSA, equal amount of HA-NRF2 and K-Glu NRF2 were subjected to binding with ARE-Biotin labelled DNA probe (NRF2(ARE) EMSA Kit, Signosis) according to manufacturer instructions.

### RNA-Seq data analysis

Illumina Truseq adapter, polyA, and polyT sequences were trimmed with cutadapt v2.3 using parameters “cutadapt -j 4 -m 20 --interleaved -a AGATCGGAAGAGCACACGTCTGAACTCCAGTCAC -A AGATCGGAAGAGCGTCGTGTAGGGAAAGAGTGT Fastq1 Fastq2 | cutadapt --interleaved -j 4 -m 20 -a “A^48^” -A “A^48^” – | cutadapt -j 4 -m 20 -a “T^48^” -A “T^48^” -”. Trimmed reads were aligned to human genome version 38 (hg38) using STAR aligner v2.7.0d 0221 ^55^ with parameters according to ENCODE long RNA-seq pipeline (https://github.com/ENCODE-DCC/long-rna-seq-pipeline). Gene expression levels were quantified using RSEM v1.3.1^56^. Ensembl gene annotation version 84 was used in the alignment and quantification steps. RNA-seq sequence, alignment, and quantification quality was assessed using FastQC v0.11.5 (https://www.bioinformatics.babraham.ac.uk/projects/fastqc/) and MultiQC v1.8^57^. Biological replicate concordance was assessed using principal component analysis (PCA) and pair-wise pearson correlation analysis. Lowly expressed genes were filtered out by applying the following criterion: estimated counts (from RSEM) ? number of samples * 5. Filtered estimated read counts from RSEM were compared using the R Bioconductor package DESeq2 v1.22.2 based on generalized linear model and negative binomial distribution^58^. Genes with Benjamini-Hochberg corrected p-value < 0.05 and fold change ≥ 2.0 or ≤ −2.0 were selected as differentially expressed genes. Differentially expressed genes were analyzed using Ingenuity Pathway Analysis (Qiagen, Redwood City, USA). RNA-seq main and supplemental figures were plotted using ggplot2 (H. Wickham. ggplot2: Elegant Graphics for Data Analysis. Springer-Verlag New York, 2016) and ComplexHeatmap^59^.

### TCGA survival analysis

Gene expression (RNA-seq) and clinical data from TCGA Pan-Cancer 2018^60^ were downloaded from cBioPortal^61^. Survival analysis was performed in R version 4.0.2 using survival (Therneau T (2020). A Package for Survival Analysis in R. R package version 3.2-7, URL: https://CRAN.R-project.org/package=survival.), survminer (Alboukadel Kassambara, Marcin Kosinski and Przemyslaw Biecek (2020). survminer: Drawing Survival Curves using ‘ggplot2’. R package version 0.4.8.999. http://www.sthda.com/english/rpkgs/survminer/), and maxstat (Torsten Hothorn (2017). maxstat: Maximally Selected Rank Statistics. R package version 0.7-25. https://CRAN.R-project.org/package=maxstat) packages. Optimal cutpoint for the categorization of TCGA samples as ‘high’ and ‘low’ GCDH expressors in each cancer type was determined using surv_cutpoint() and surv_categorize() functions from survminer package.

### Measurement of cellular respiration

siRNA transfected A375 cells were plated at a density of 10,000 per well in a Seahorse XFp culture plate and cultured overnight before changing the medium to Seahorse XF base medium containing 1 g/l glucose and 2 mM glutamine, pH 7.4, and assaying oxygen consumption and extracellular acidification rates in the Seahorse XFp with successive additions of 1.5 mM oligomycin and 1.0 mM FCCP.

### Statistical Analysis

Statistical significance between two groups was assessed by the unpaired Student’s t-test. Ordinary one-way ANOVA was used to analyze more than two groups. Two-way ANOVA was utilized to analyze cell proliferation at multiple timepoints. GraphPad Prism 5 software Graphpad 8.0.0 (224), La Jolla, CA) was used for to perform all statistical calculations.

## Acknowledgements

We thank members of the Prebys Center for Chemical Genomics for help with the screen for GCDH inhibitors and members of the Ronai lab for discussions. NCI support (R35CA197465) to ZR and SBP support for translational initiatives (to ES and ZAR) are gratefully acknowledged. Sanford Burnham Prebys Shared Resources are supported by an NCI Cancer Center Support Grant (P30 CA030199).

## Author contributions

SV, GP, AK, and YF performed experiments; DC, ER, and RM performed bioinformatic analyses;; ZAR, SV, and ES designed the studies, and ZAR, SV, and ES wrote the manuscript.

## Supplementary Figure Legends

**Figure SI.**
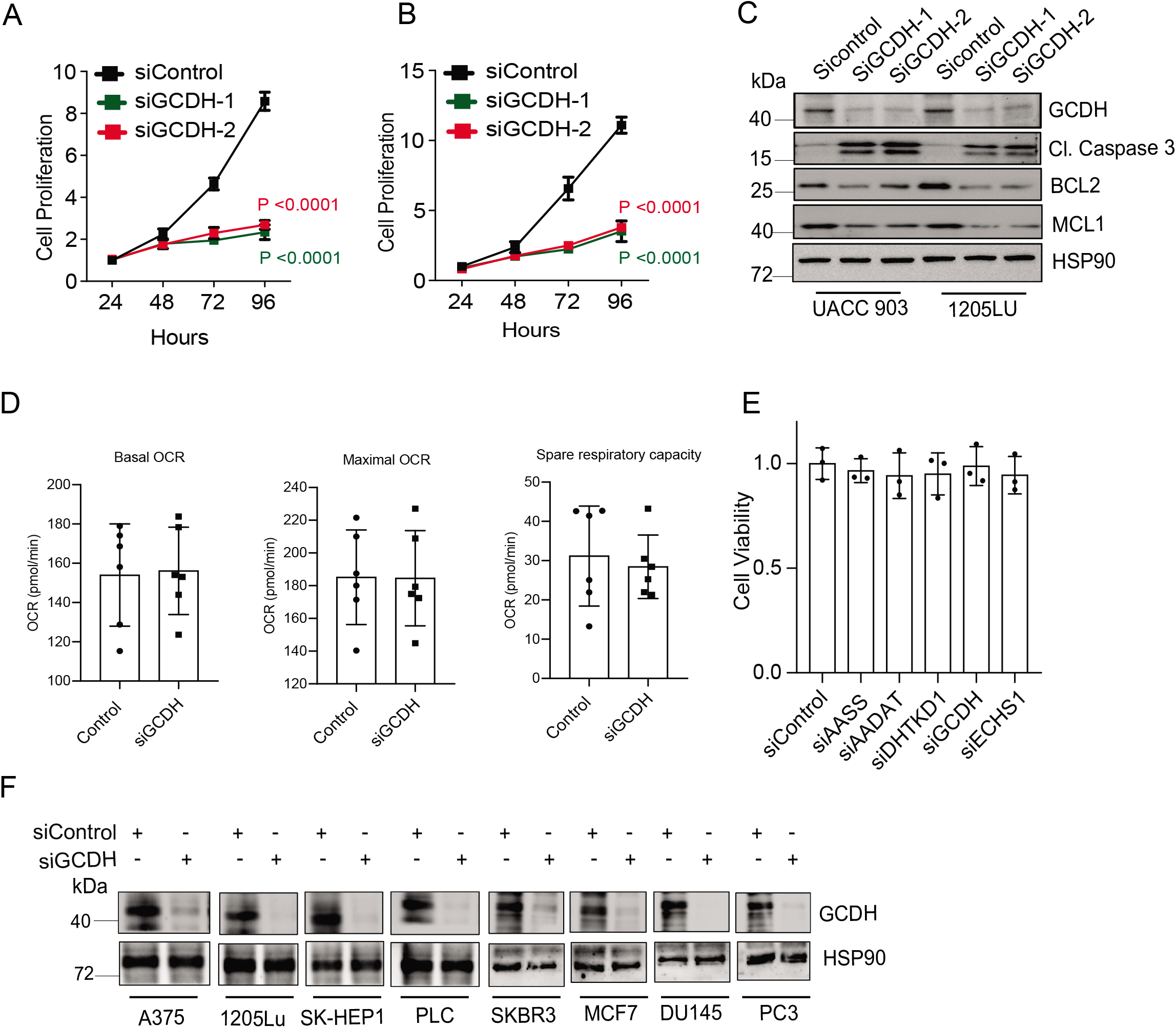
GCDH inhibition promotes apoptosis in melanoma cells. (A, B) Cell growth upon GCDH knock down using two independent siRNAs for 0-96 hr in UACC903 (A) or 1205LU (B). Cell growth was analyzed by cell counting at indicated time points. (C) UACC903 or 1205LU cells were transfected with siRNA against GCDH for 72 hours, and western blot analysis was done using indicated antibodies. (D) Measurement of basal, maximum oxygen consumption rate (OCR) and spare respiratory capacity by Agilent Seahorse XF Analyzers. siRNA targeting GCDH was transfected into A375 cells before analysis. (E) Cell viability assay of immortalized H3A cells, 96 hr after transfection with indicated constructs. Cell viability was measured by quantifying crystal violet staining. Data are presented as the mean ± SEM. Statistical significance (indicated p value or ns-not significant w.r.t control) was calculated using unpaired *t*-test.

**Figure S2.**
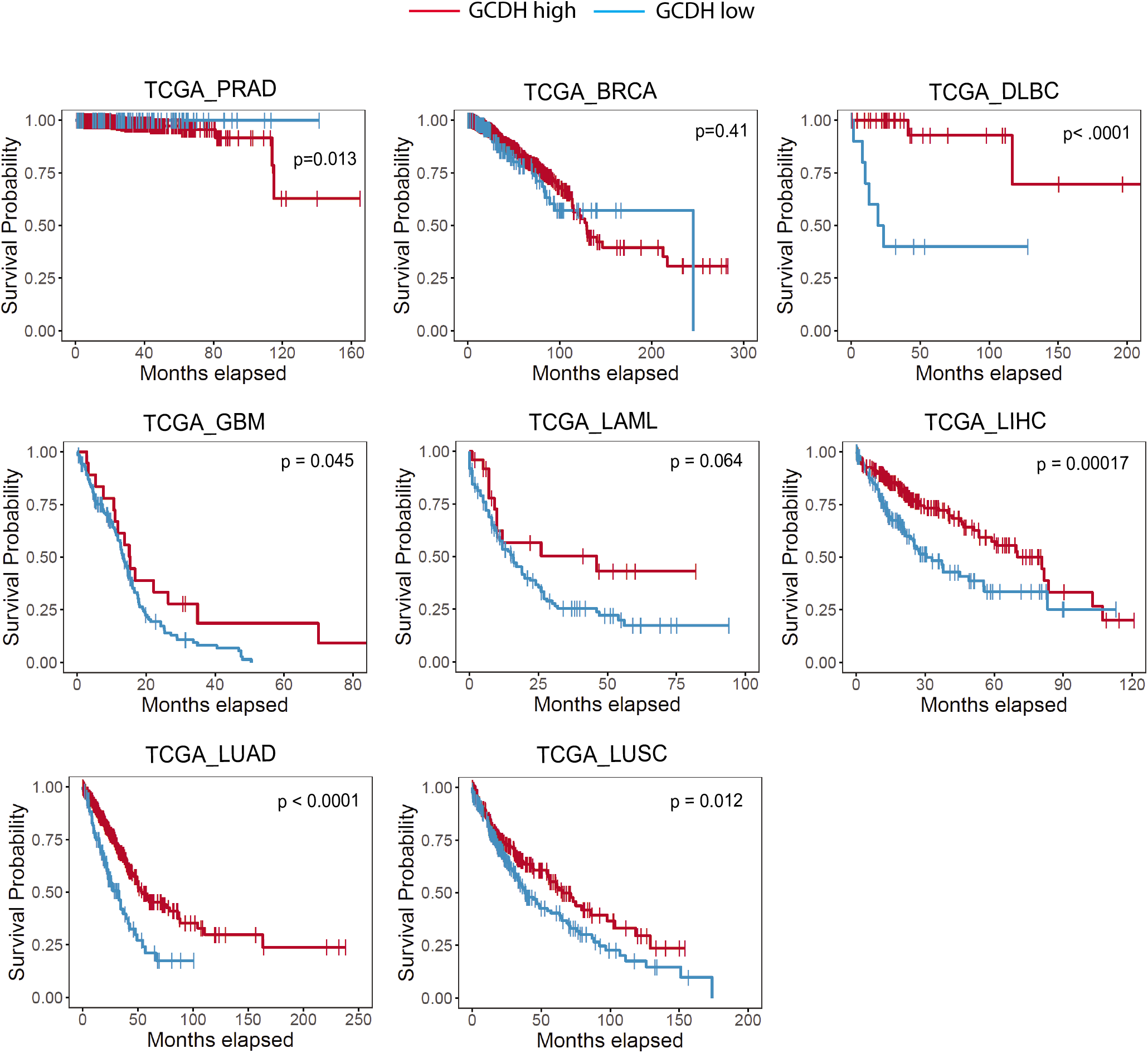
GCDH expression coincides with patient outcome in melanoma. Survival correlation analysis of GCDH expression in various cancer subtypes (prostate adenocarcinoma (PRAD), breast cancer (BRCA), diffuse large B-cell lymphoma (DLBC), glioblastoma (GBM), acute myeloid leukemia (LAML), liver hepatocellular carcinoma (LIHC), lung adenocarcinoma (LUAD), lung squamous cell carcinoma (LUSC) using TCGA.

**Figure S3.**
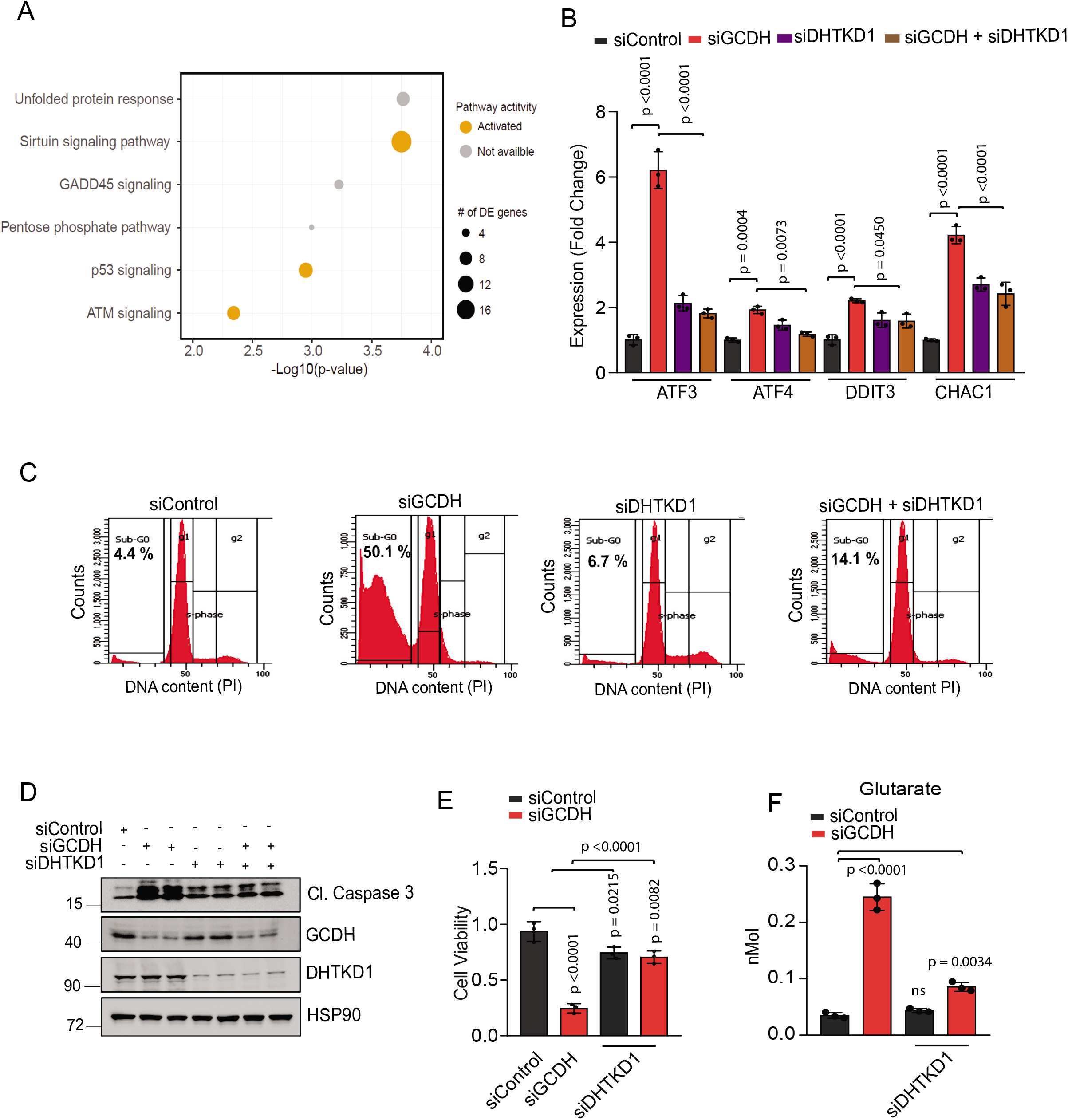
DHTKD1 inhibition rescues gene expression changes and cell death phenotypes seen following GCDH inhibition. (A) Gene set enrichment analysis to identify signaling pathway affected upon GCDH KD in A375 cells identified by RNA-seq analysis. (B)RT-qPCR analysis in UACC903 cells for relative expression of *ATF3, ATF4, DDIT3* and *CHAC1* following GCDH-KD, DHTKD1-KD alone or GCDH-DHTKD1 double KD. (C) SubG0 DNA content analysis by flow cytometry to measure apoptosis in A375 cells. A375 cells were transfected with indicated siRNAs for 72 hours and then harvested for fixation in ethanol and staining with propidium iodide (PI). (D) Western blot analysis to measure GCDH, DHTKD1 and Cl. caspase 3 protein levels in A375 lines following GCDH-KD, DHTKD1-KD alone or GCDH-DHTKD1 double KD. (E) Rescue of cell death in GCDH-KD 1205LU upon DHTKD1-KD. Cell viability was measured by quantifying crystal violet staining. (F) GC-MS analysis to measure glutarate concentrations in A375 cells. Data are presented as the mean ± SEM. Statistical significance (indicated p value or ns-not significant w.r.t control) was calculated using unpaired *t*-test except for (F), (E) and two-way ANOVA for (B) and (E).

**Figure S4.**
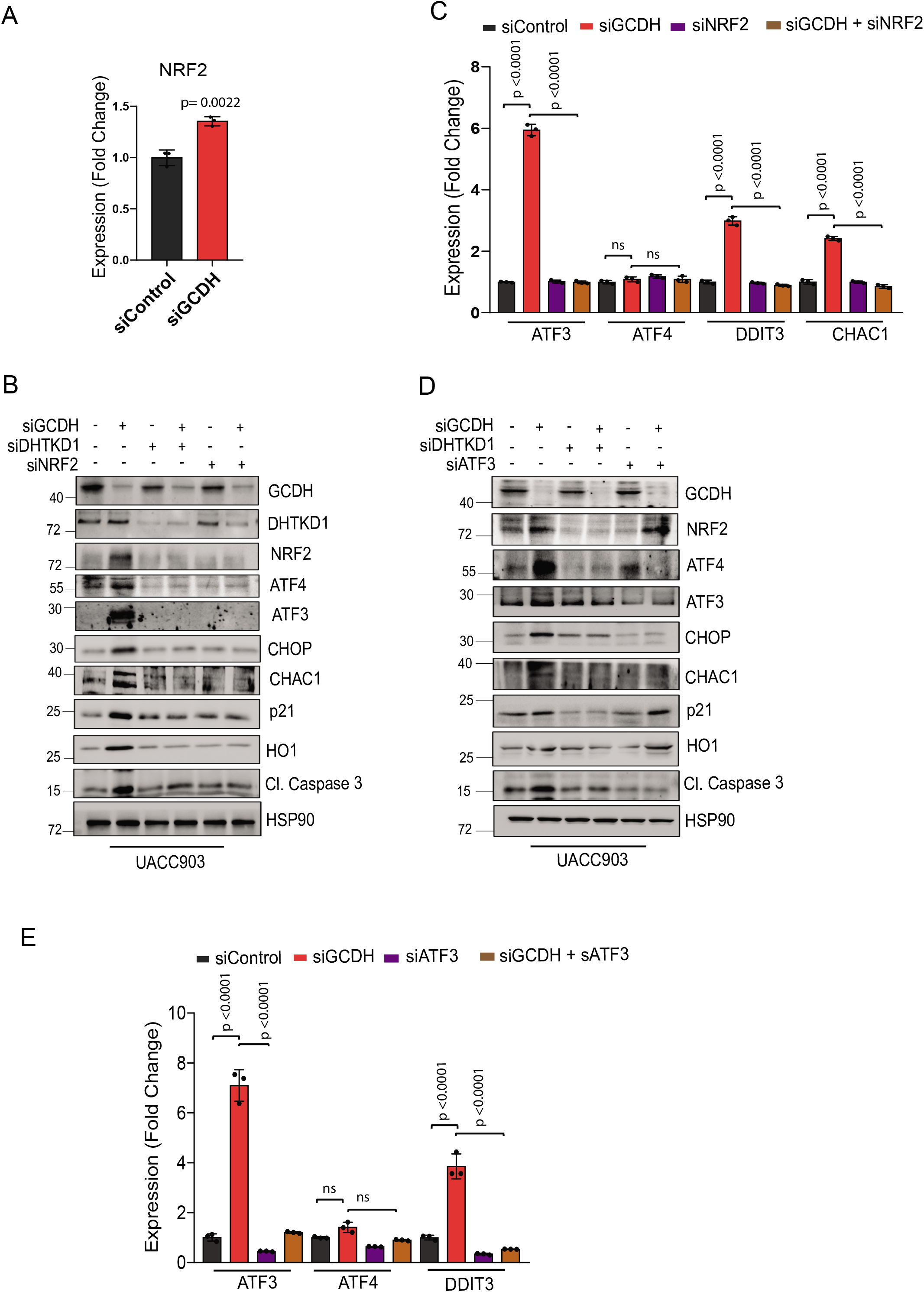
GCDH activity in melanoma cells antagonizes NRF2-mediated activation of ATF3/4 downstream apoptotic signaling. (A) RT-qPCR analysis of NRF2 mRNA expression upon GCDH KD in A375 cells. (B) Western blot analysis of indicated proteins in UACC903 cells, 72 hr following transfection with various siRNAs. (C) RT-qPCR analysis of *ATF3, ATF4, DDIT3,* and *CHAC1* expression levels in UACC903 cells 72 hr following transfection with indicated siRNAs. (D) Western blot analysis of indicated proteins following in UACC903 cells 72 hr following transfection with siRNAs. (E) RT-qPCR analysis of *ATF3, ATF4, DDIT3,* and *CHAC1* expression levels in UACC903 following transfection with indicated siRNAs. Data are presented as the mean ± SEM. Statistical significance (indicated p value or ns-not significant w.r.t control) was calculated using unpaired *t*-test except for (A), and two-way ANOVA for (C) and (E).

**Figure S5.**
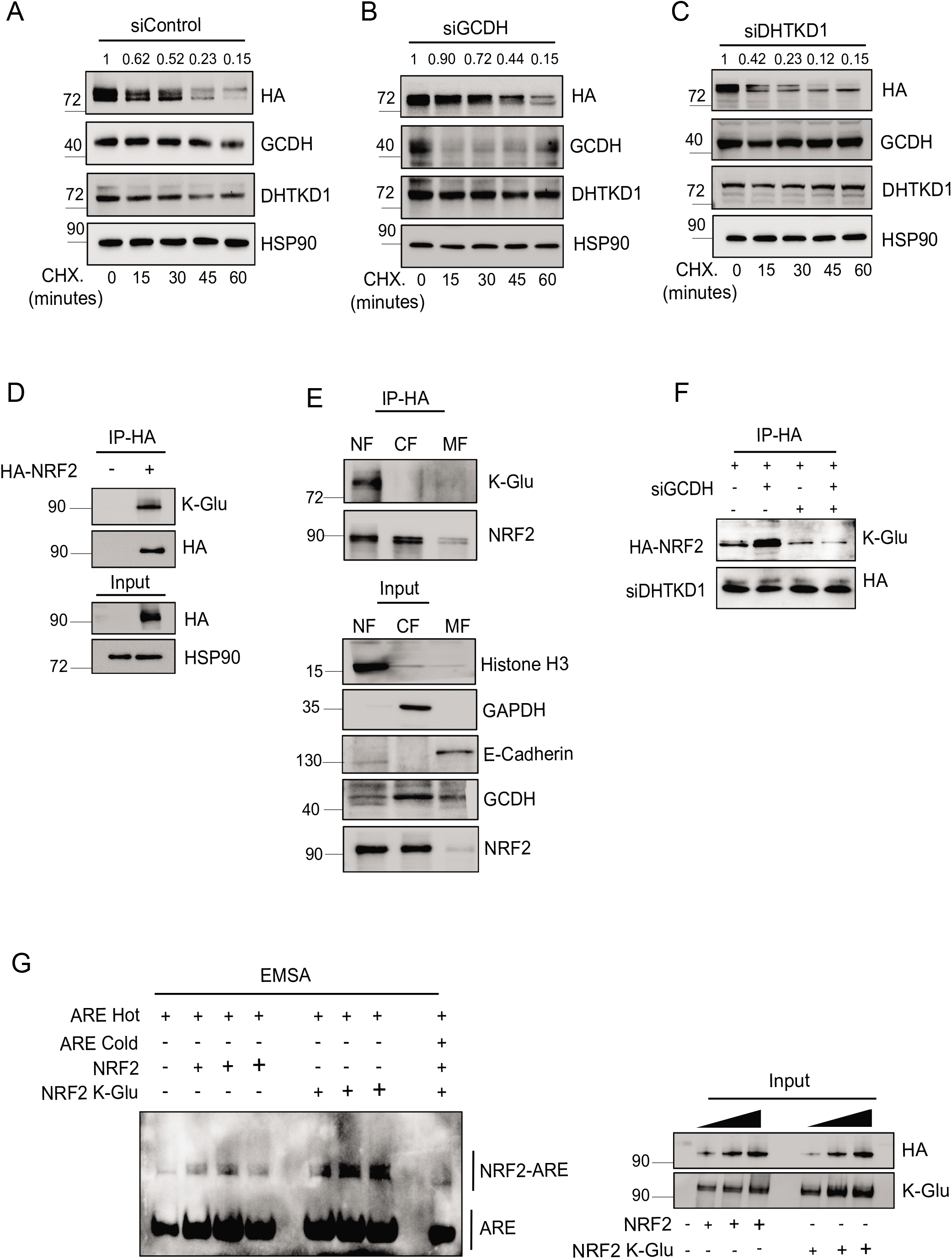
Lysine glutarylation increases NRF2 stability and antagonizes KEAP1 binding. (A, B) Cycloheximide (CHX) chase analysis to measure HA-NRF2 stability in control and (B) GCDH KD and (C) DHTKD1-KD HEK-293T cells ectopically expressing HA-NRF2. HEK293T cells were transfected with indicated constructs for 72 hours and then treated with 10 μM Cycloheximide (CHX) for different time followed by western blotting with indicated antibodies. After quantification, the signals obtained in panel A, B and C were used to calculate the HA-NRF2/HSP90 ratios and described with respect to CHX. treatment period. (D) Immunoprecipitation and Western blot analysis of HA-NRF2 from HEK293T transfected with indicated constructs. HEK 293T cells ectopically expressing HA-NRF2 and treated with the 10 μM proteasomal inhibitor MG132 followed by IP/Western blotting analysis with antibodies to detect K-Glu PTM and HA-NRF2. (E) Enrichment of NRF2 glutarylation in nuclear fraction was measured by HA-NRF2 pull-downs from HEK293T cells transfected with HA-NRF2, after an initial cell fractionation step using MF-membrane fraction; CF-cytoplasmic fraction; NF-nuclear fraction. Successful cell fractionation was confirmed by immunoblotting for specific markers of MF(E-cadherine), CF (GAPDH), and NF (Histone H3). (F) Relative DNA binding activity of purified HA-NRF2 or K-Glu-NRF2 were measured by in vitro NRF2/ARE EMSA (Electrophoretic-Mobility Shift Assay) using native (non-denaturing) polyacrylamide gels electrophoresis followed by western blotting. A representative image of n = 3 independent experiments is shown.

